# Confirming Multiplex Q-PCR Use in COVID-19 with Next Generation Sequencing: Strategies for Epidemiological Advantage

**DOI:** 10.1101/2022.02.22.481485

**Authors:** Rob E. Carpenter, Vaibhav Tamrakar, Harendra Chahar, Tyler Vine, Rahul Sharma

## Abstract

Rapid classification and tracking of emerging SARS-CoV-2 variants are critical for understanding the transmission dynamics and developing strategies for interrupting the transmission chain. Next-Generation Sequencing (NGS) is an exceptional tool for whole-genome analysis and deciphering new mutations. The technique has been instrumental in identifying the Variants of Concern and tracking this pandemic. However, NGS remains expensive and time-consuming for large-scale monitoring of COVID-19. This study analyzed a total of 78 de-identified samples that screened positive for SARS-CoV-2 from two timeframes, August 2020 and July 2021. All 78 samples were classified into WHO lineages by whole genome sequencing then compared with two commercially available Q-PCR assays for spike protein mutation(s). The data showed good concordance with Q-PCR and NGS analysis for specific SARS-COV-2 lineages and characteristic mutations.

Deployment of Q-PCR testing to detect known SARS-COV-2 variants may be extremely beneficial. These assays are quick and cost-effective, thus can be implemented as an alternative to sequencing for screening known mutations of SARS-COV-2 for clinical and epidemiological interest. The findings support the great potential for Q-PCR to be an effective strategy offering several COVID-19 epidemiological advantages.

## Introduction

The coronavirus (COVID-19) pandemic started in December 2019 in Wuhan, China. It has been considered one of the deadliest infectious disease outbreaks in recent world history. The causative agent of COVID-19 is the Severe Acute Respiratory Syndrome Coronavirus 2 (SARS-CoV-2). SARS-CoV-2 is a positive-sense RNA virus belonging to the coronaviridae family, genus betacoronavirus, and subgenus arbovirus.^1,2^ COVID-19 has had devastating effects on the human population and to date is estimated to have caused over 5 million deaths worldwide.^3^ Rapid accurate diagnosis of SARS-CoV-2 is the most crucial step in the management of COVID-19—mostly achieved with gene amplification by reverse transcription polymerase chain reaction (RT–PCR). The assays detect highly conserved regions in the open reading frame (ORF) 1a or 1b, and nucleocapsid (N) gene of SARS-CoV-2.^4–6^

Currently, the virus continues to be a global agent of infection. The highly mutagenic nature of SARS-CoV-2 assaults many countries with second or third waves of the outbreak.^7,8^ Mutations with higher transmissibility, a more intense disease state, and that are less likely to respond to vaccines or treatments, have been classified by the World Health Organization (WHO) as Variants of Concern (VOC; Table 1). Recent epidemiological reports released by WHO indicated four VOC: 1) Alpha (B.1.1.7), first reported in the United Kingdom (UK) in December 2020; 2); Beta (B.1.351), first reported in South Africa December 2020; 3) Gamma (P.1), first reported in Brazil January 2021; and 4) Delta (B.1.617.2), first reported in India December 2020.^7^ The virus has demonstrated that genomic changes in its receptor binding domain (RBD)—a region of the spike protein that studs SARS-CoV-2 to the outer cell surface—inserts increased capacity to strike in several outbreak phases in different parts of the world.^9^ More recently, South Africa reported a new SARS-CoV-2 variant to the WHO. Omicron (B.1.1.529) was first detected in specimens collected in Botswana. On November 26, 2021, the Technical Advisory Group on SARS-CoV-2 Virus Evolution (TAG-VE) advised WHO to designate B.1.1.529 as the fifth VOC.^10^ Genomic mutations for classifying the VOCs are summarized in Table 1.

**Table 1:** World Health Organization (WHO) Designated Variants of Concern (VOC)^11^

| WHO Label | Alpha | Beta | Gamma | Delta | Omicron |  |
| --- | --- | --- | --- | --- | --- | --- |
| Pango Lineage | B.1.1.7 | B.1.351 | P.1 | B.1.617.2 | B.1.1.529 |  |
| Classifying Mutation(s) | Δ69/70 | K417N | K417N/T | T19R | A67V | E484A |
|  | Δ144Y | E484K | E484K | (G142D*) | Δ69-70 | Q493K |
|  | (E484K*) | N501Y | N501Y | Δ156 | T95I | G496S |
|  | (S494P*) | D614G | D614G | Δ157 | G142D | Q498R |
|  | N501Y |  |  | R158G | Δ143-145 | N501Y |
|  | A570D |  |  | L452R | Δ211 | Y505H |
|  | D614G |  |  | T478K | L212I | T547K |
|  | P681H |  |  | D614G | ins214EPE | D614G |
|  |  |  |  | P681R | G339D | H655Y |
|  |  |  |  | D950N | S371L | N679K |
|  |  |  |  |  | S373P | P681H |
|  |  |  |  |  | S375F | N764K |
|  |  |  |  |  | K417N | D796Y |
|  |  |  |  |  | N440K | N856K |
|  |  |  |  |  | G446S | Q954H |
|  |  |  |  |  | S477N | N969K |
|  |  |  |  |  | T478K | L981F |

\*Detected in some sequences but not all.

There continues to be a need for swift and cost-effective SARS CoV-2 variant detection and monitoring. Genomic sequencing is the gold standard and most reliable method for the detection of such changes in the viral genome. The standard Sanger sequencing method^12^ is highly accurate but it can only sequence a small fraction of the genome. Sanger sequencing is also laborious, time-consuming, and expensive for large-scale sequencing projects that require rapid turnaround times. These attributes make Sanger sequencing less attractive for SARS CoV-2 sequencing for variant identification and monitoring.

Targeted Next-Generation Sequencing (NGS) is also a reliable method to identify variant strains of pathogens, including viruses.^13^ The principal advantage of NGS over other techniques like Sanger sequencing or RT-PCR is that scientists and laboratorians do not require prior knowledge of existing nucleotide sequences. Moreover, NGS has higher discovery power and higher throughput.^13^ In the current pandemic, NGS has widely been employed to detect and identify novel mutated viral variants of SARS COV-2.^14^ Although widespread adoption of NGS in clinical laboratories offers effective variant discovery, several challenges impede the routine use of NGS in these settings. Besides the need for multifaceted NGS validation studies,^15^ NGS testing is complicated by the high level of necessary human expertise and the cost-scalability for routine pathogen detection. Moreover, the interpretation of results generated by NGS can be intricately complex and their applicability to clinical decision making is another issue altogether. These complexities pose the need to progress practical methodologies to identify SARS COV-2 mutagenic variants quickly and cost-effectively. The current study proposes that RT–PCR techniques could be utilized to extend the detection of known mutation(s) by designing primers and probes against the previously identified loci of targeted SARS COV-2 mutations evidence by NGS.

## Methodology

This study used a total of 78 de-identified sample remnants from nasopharyngeal or oropharyngeal swabs (catalog# 202003, Nest Biotechnology Jiangsu, China) collected from patients that screened positive for the presence of SARS-CoV-2 following RNA extraction and RT-PCR at Advanta Analytical Laboratories (https://aalabs.com/) in Tyler, Texas. All the clinical samples were collected from Texas residents who first tested positive for SARS-Cov-2 with established protocols targeting N1 and N2 genes with established primer and probe design (see Appendix A1 for design). Eleven samples were collected and archived during the early (August 2020) pandemic. The remaining 67 samples were collected in July 2021 following the global outbreak of the Delta (B.1.617.2) variant. We qualified the samples with moderate to high viral amplification by cycle threshold (Ct) values ≤30 for N1 and N2 genes following RT-PCR testing on the LightCycler® 480 System (Roche). We also included 4 samples with low viral amplification (30<Ct>35; PreD 15, 16, 18, and 19) in the study to evaluate the applicability of RT-PCR and NGS for reduced Ct values (Ct values are inversely proportional to amplification thresholds; Table 2).

**Table 2:**
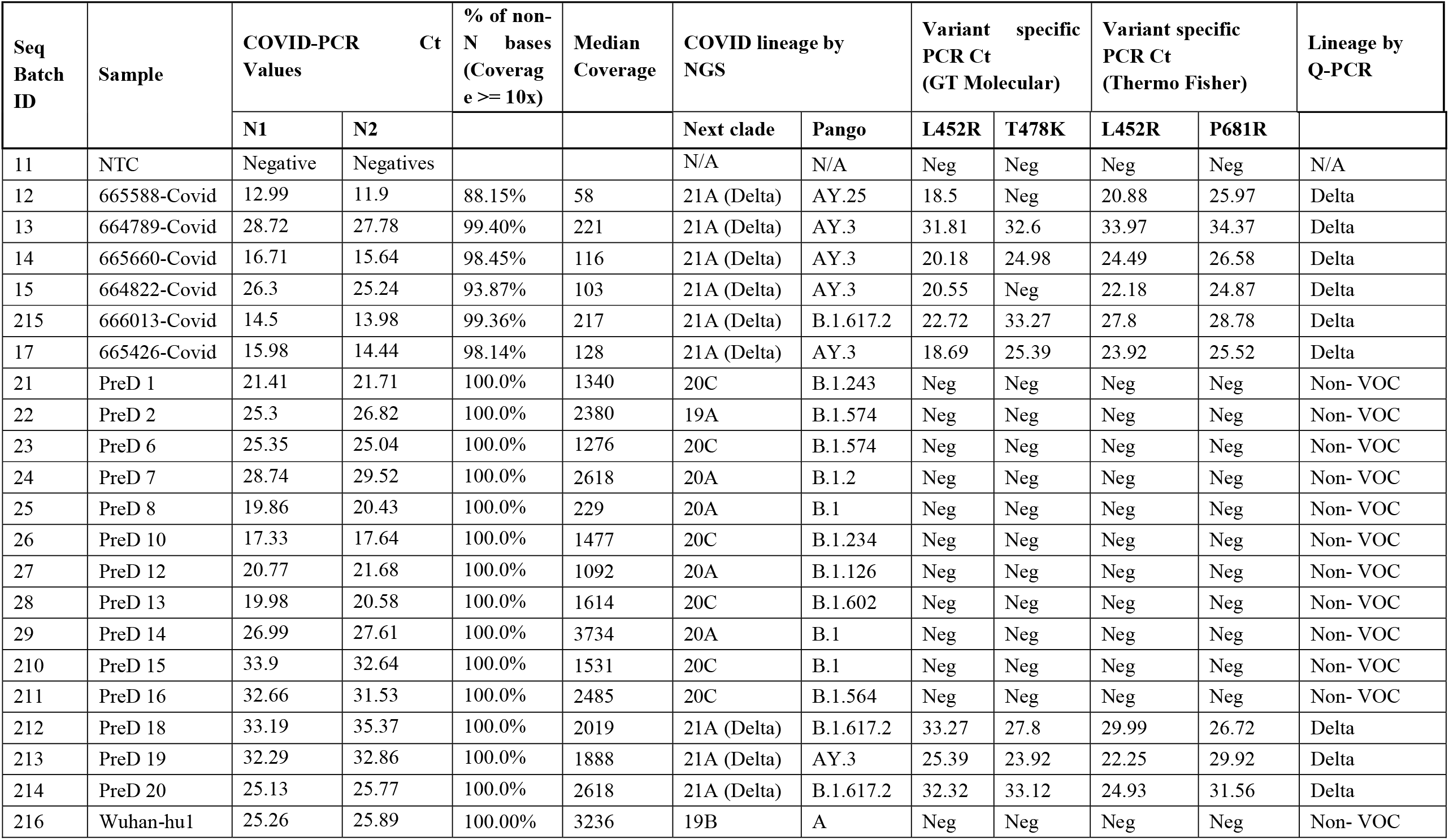
Data of representative samples (*n*=20) sequenced and classified into Delta and non-VOC variants using NGS and Q-PCR assays

Next, library preparation and NGS was performed on each sample and analyzed for lineage and variant detection. We obtained and reviewed the whole-genome sequence of the samples in two phases. A total of 64 samples were sequenced at Fulgent Genetics (https://www.fulgentgenetics.com/) and 20 samples were sequenced at Advanta Analytical Laboratories (there was an overlap of 6 samples sequenced by both laboratories). After NGS results were verified, we incorporated phylogenic analysis aimed at investigating the conservation of spike protein in reference sequences vs clinical strains of SARS-CoV-2 from our study using bioinformatics tools. Once NGS lineage and variant detection was established, we progressed to SARS CoV-2 lineage assignment and variant detection using qualitative polymerase chain reaction (Q-PCR) for comparison. We approached this by testing all 78 samples with two commercially available PCR assays from two vendors (GT molecular biology [Colorado USA], and Thermo Fisher Scientific [Massachusetts, USA]). Results were analyzed and linked to the NGS-based variant detection of the same samples (see Appendix A2 for expanded description of RNA extraction, library preparation and sequencing, NGS data analysis, phylogenic analysis, and SARS CoV-2 lineage assignment using Q-PCR).

## Results

The 78 randomly selected positive SARS COV-2 samples were from two separate periods in the pandemic. NGS of the 11 samples from August 2020 revealed eight different lineages, but none of the lineages were VOC according to the WHO classification. All samples (100% [67/67]) sequenced from July 2020 revealed the SARS COV-2 Delta (B.1.617.2, AY3, and AY 25) VOC with sub-lineages AY.1 to AY.3. Incidentally, the six samples concurrently sequenced at both laboratories were identified as Delta (B.1.617.2) VOC. Unfortunately, raw data (FASTQ files) were not available from the samples sequenced at Fulgent Genetics. However, the raw data from the 20 samples sequenced at Advanta Analytical Laboratories was analyzed for phylogenetic relationship and mutation discovery (Table 2). This data revealed novel mutations belonging to existing prominent lineages along with convergent mutations of different lineages and one unique mutation (1).

We then turned our focus to testing the 67 Delta (B.1.617.2) samples by using Q-PCR methodology targeting three (L452R, T478K, &P681R) characteristic mutations identified through sequencing. We tested each sample using two different commercially available (Thermo Fisher and GT Molecular) assays and compared the results. The Delta (B.1.617.2) classifying mutation (L452R) was correctly identified by GT Molecular Q-PCR based assay, and the test showed 100% concordance for all 67 samples that were sequenced as Delta (B.1.617.2). However, the Thermo Fisher assay for the same target (L452R) did not amplify in 4 out of 67 samples otherwise identified as Delta variant by NGS (Table 3; Supplementary Table-S1). Overall, GT Molecular assay targeting the L452R mutations had 4.21 ± 2.3 lower Ct value when the same RNA template was tested with both assays suggesting higher sensitivity of the GT Molecular assay. Moreover, 5 of 67 samples were negative for T478K (GT Molecular), and 12/67 were negative for P681R specific PCR (Thermo Fisher) using Q-PCR, (Table 3). Unfortunately, we could not verify the absence of these mutations because NGS data was not available for the 64 samples sequenced at Fulgent Genetics. Thus, the L452R mutation remained the most informative marker for Q-PCR based detection of the Delta variant. All 11 samples sequenced as non-Delta variants were negative for all three Delta variant-specific mutations (Table 3). Interestingly, a Beta and Gamma variant classifying mutation (E484K) was identified (both by Q-PCR assays and NGS) in one sample, which is otherwise classified as Delta variant by NGS and carries a L452R mutation. This mutation combination is suggestive of the continuous evolution of the coronavirus genome.

**Table 3:**
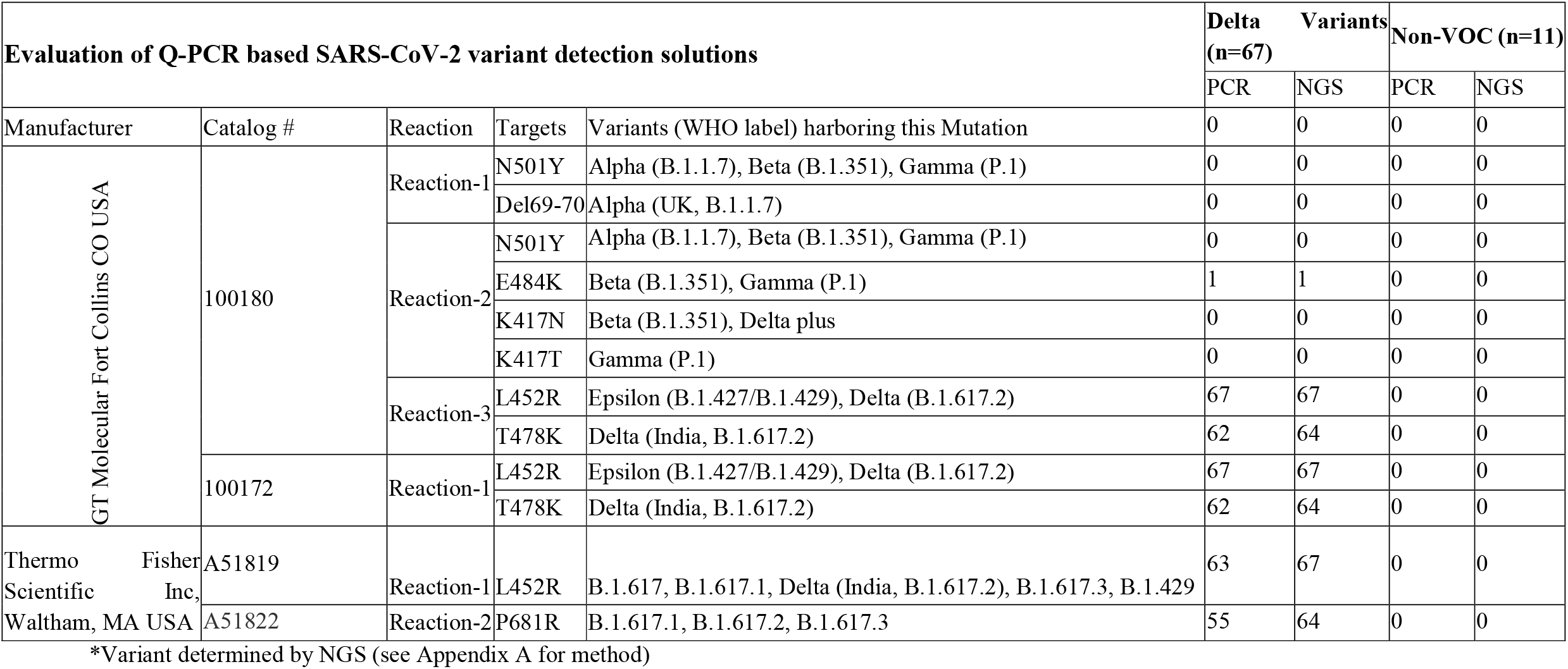
Comparative detection of VOC classifying mutations with RT-PCR and NGS based approaches

Of note, the 4 samples with lower viral amplification (35<Ct>30) that were included in this study were able to be characterize by NGS and both Q-PCR assays. Two out of the four samples were identified as Delta (B.1.617.2) variants with the remaining two identified as non-VOC. Therefore, NGS and Q-PCR methodologies can potentially be used for SARS CoV-2 variant detection from the samples with lower viral amplification (1000-10 copies).

In addition to NGS and Q-PCR variant concordance of the 78 samples, the results of this study reveal significantly reduced diversity of SARS COV-2 variants from July 2020 to August 2021. We detected 8 lineages among the 11 samples tested from July 2020 compared to a single Delta variant lineage with three sub-lineages (Delta) among 67 samples collected in August 2021 (Figure 2). These findings are important for understanding the evolution of SARS COV-2 variants in Texas (Figure 2) and support other studies showing predominance and infectivity of the Delta variant.^16^

**Figure 1:**
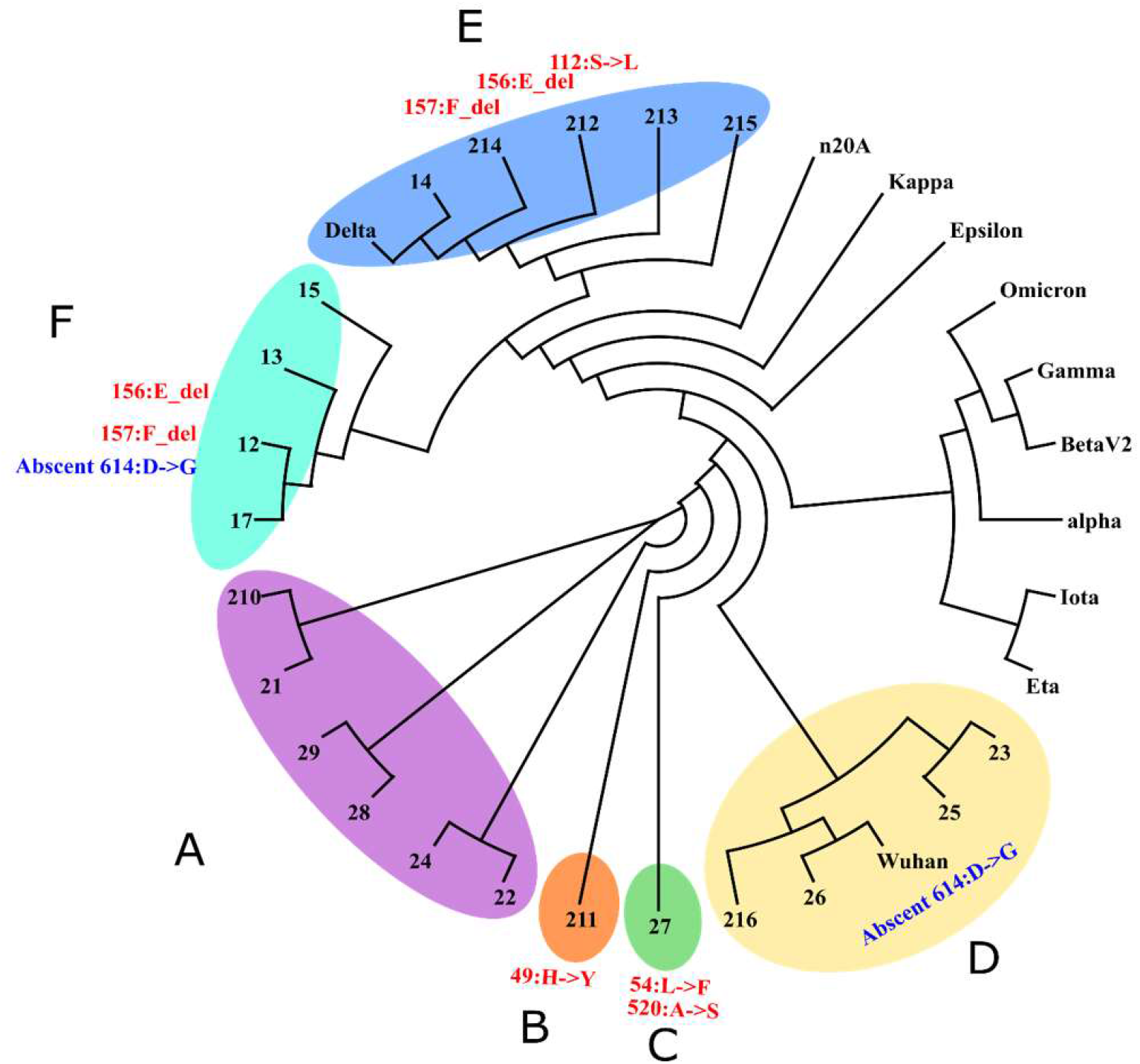
Minimum spanning tree of the SARS COV-2 variants identified in samples (*n*=20) collected August 2020 and July 2021. *Note:* No Brazilian or United Kingdom lineages were identified. Two groups of samples (F & D) lacked omnipresent mutation (614:D->G) which is present in most variants of concern. Group F is of particular interest as it had most of the Delta variant mutations except 614:D->G which is present in all the samples unless otherwise marked. Group E on the other hand is Delta lineage with a novel mutation among them (i.e. 112:S->L). Group A was identical to the Wuhan strain except for 614:D->G mutation. Two samples are carrying unique lineage and mutation sets: Set B had a single sample with mutation 49:H->Y which is novel and not found in any other lineage and Set C also had a single sample with unreported mutation pair 54:L->F and 520:A->S.

**Figure 2:**
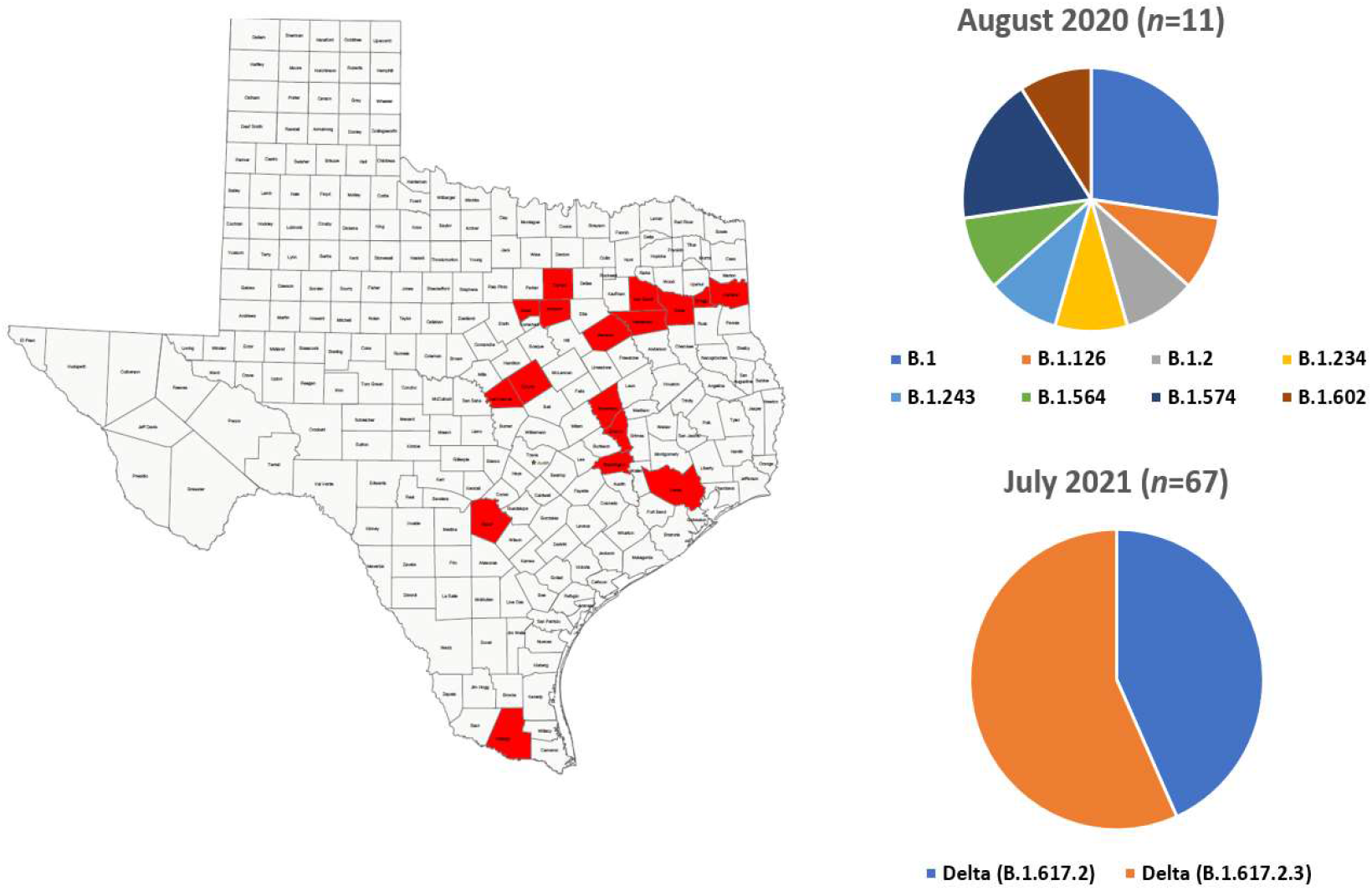
Evolution of the SARS COV-2 variant in Texas over one year (August 2020 to July 2021).

## Discussion

The emergence of new SARS COV-2 variants with higher infection rates and morbidity continues to cause the global scientific community concern. To manage further transmission and control of infection, genomic surveillance is important for the identification and tracking of novel variants. NGS is a very useful tool for identifying new strains of COVID-19 and other infectious pathogens. NGS can be used to detect novel pathogen mutations and can also be used to determine the rate of pathogen evolution. Although the unavailability of the raw data (FASTQ files) from the 67 samples remains the limitation of the study, phylogenetic analysis of the 20 samples tested at Advanta Analytical Laboratories were clustered as expected; all the VOC and Non-VOC samples were grouped appropriately.

Although NGS is the most reliable method for detecting mutations in SARS-CoV-2, the methodology is not practically applicable for large-scale surveillance, particularly in resource-limited settings. Factors like continuous validation studies, logistic challenges, database validity, cost-benefit analysis, and high technical expertise make the implementation of NGS in routine clinical settings difficult. Comparatively, Q-PCR—a gold standard for diagnosing SARS-CoV-2—is a method that can be extended for variant detection and monitoring in clinical settings.

The novelty of this research is that it demonstrated that Q-PCR is as effective as NGS in detecting SARS-CoV-2 mutations. Two Q-PCR-based assays for the detection of SARS-CoV-2 mutagenic variants were tested and compared with NGS data. Both assays were able to detect L452R mutation with 100% (67/67; GT Molecular) and 94% (63/67; Thermo Fisher) accuracy when compared to NGS. While NGS is an essential tool for sequencing the entire genome and identification of new mutations, this study suggests Q-PCR can aptly serve as an easy to deploy, cost-effective, and time-sensitive solution for the detection of known mutations for mass surveillance. Likewise, this approach has been previously applied for surveillance of leprosy and identification of zoonotic transmission in the United States.^17,18^ The authors used NGS data to develop an algorithm for the classification of global variants and deployed Q-PCR to understand the local transmission dynamics.

The results in this study are promising because the Q-PCR lineage classification showed no mismatches when compared with the 20 sequenced samples that had raw data. Although these results are encouraging for cost-scalability of SARS CoV-2 mutation detection with Q-PCR, other research using similar virus sequencing comparison methods have been less successful. Khan and Cheung^19^ noted the presence of mismatches when comparing SARS-CoV-2 between PCR and sequencing data. Elaswad and Fawzy^20^ also found this to be the case when comparing PCR assays with available SARS-CoV-2 genomes isolated from animals. Similarly, Mai et al^21^ noted missed detections with PCR assays for influenza A (H1) when compared with sequencing. Although these studies add some concern, it does appear strategic deployment of both NGS and Q-PCR based solutions for discovery and monitoring of emerging SARS-CoV-2 mutations is likely to advance better strategies for epidemiological characteristics.

## Conclusion

There are two important take aways from this research. First, the NGS data provided further evidence of upcoming resident SARS-COV-2 lineages and suggests the continued threat of COVID-19. This finding is consistent with other research and further supports the need for rapid, cost-effective monitoring of variant mutations. Second, the current study validates that Q-PCR-based assays have potential to be a solution for more accessible variant monitoring. The data showed concordance with Q-PCR and NGS analysis for specific SARS-COV-2 lineages and characteristic mutations. Thus, deployment of Q-PCR testing for the detection of known SARS-COV-2 variants may be extremely beneficial. These assays are quick and cost-effective, thus can be implemented as an alternative to sequencing for screening known mutations of SARS-COV-2 for clinical and epidemiological interest. In developing countries, COVID-19 diagnostic centers are mostly dependent on the regional sequencing laboratories for screening the mutations in the SARS-COV-2 clinical samples. The findings in this study depicts great potential for Q-PCR to be an effective strategy offering several epidemiological advantages.

## Appendix A1 RT-PCR Primer and Probe design

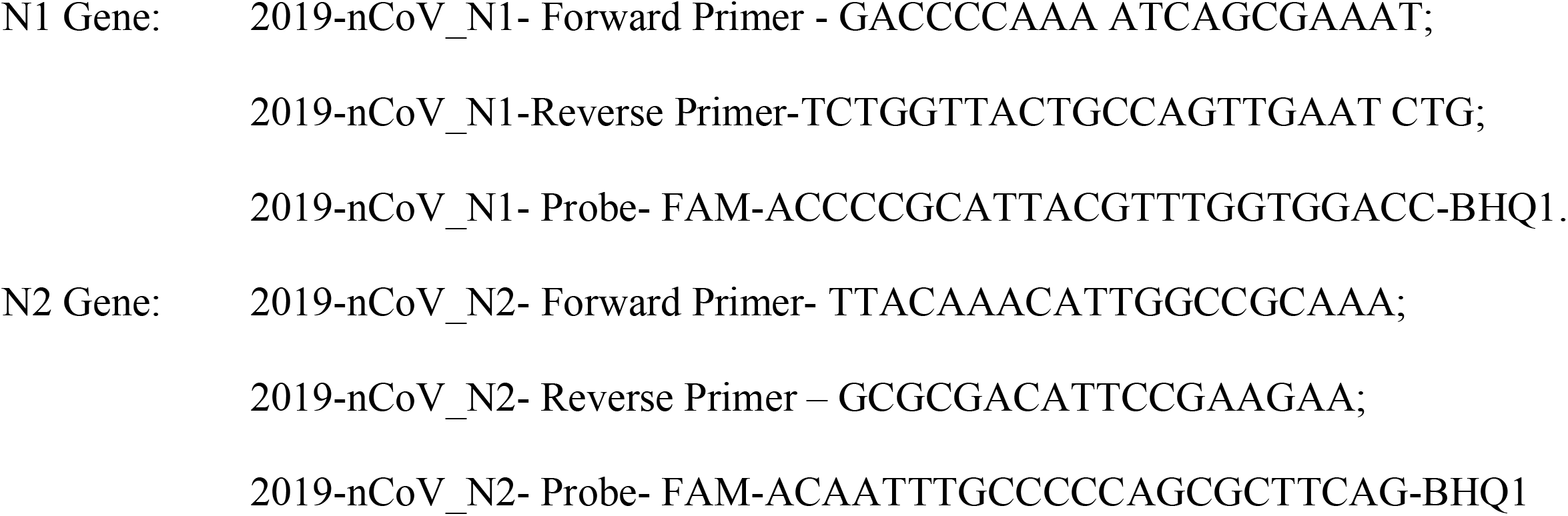

## Appendix A2

Expanded Description of RNA Extraction, Library Preparation and Sequencing, NGS Data Analysis, Phylogenic Analysis, and COVID-19 Lineage Assignment Using Q-PCR

### RNA Extraction

Each sample was extracted from nasopharyngeal or oropharyngeal swabs collected and transported to the lab in MANTACC Transport Medium or Viral Transport Medium (VTM) purchased from Criterion Clinical (https://criterionclinical.com/). RNA extraction was carried out in a pre-amplification environment within a Biosafety level 2 (BSL-2) facility. RNA isolation was performed as part of routine diagnostic testing using the Roche MagNA Pure 96 System and Viral NA Small Volume Kits. Briefly, samples were lysed with 340 uL of lysis buffer and 10 uL of proteinase K at 55°C for 10 minutes followed by extraction via the Roche MagNA Pure 96 instrument. Extracted nucleic acids were immediately sealed with a PCR clean sealing film (Cat # T329-1 Simport Scientific Inc. QC J3G 4S5 Canada) and frozen at −80°C until sequencing was imminent.

### Library Preparation and Sequencing

The libraries were prepared using Illumina COVIDSeq protocol (Illumina Inc, USA). Total RNA was primed with random hexamers and first-strand cDNA was synthesized using reverse transcriptase. The SARS-CoV-2 genome was amplified using the two sets of primers (COVIDSeq Primer Pool-1 &2) in two multiplex PCR protocols to produce two sets of amplicons spanning the entire genome of SARS-CoV-2. The PCR amplified product was then processed for tagmentation and adapter ligation using 24 IDT for Illumina Nextera UD Indexes Set A. Further enrichment and cleanup were performed as per protocols provided by the manufacturer (Illumina Inc, USA). A COVIDSeq positive control (Wuhan-Hu-1) and one no template control (NTC) was processed with each batch of libraries. Individual libraries were quantified using Qubit 2.0 fluorometer (Invitrogen, Inc.) and fragment sizes were analyzed in Agilent 5200 Fragment Analyzer. Libraries were pooled into an equimolar concentration. The library pool was further normalized to 1nM concentration and denatured and neutralized with 0.2N NaOH and 400mM Tris-HCL (pH-8) respectively. Denatured libraries were further diluted to 2 pM loading concentration. Dual indexed paired-end sequencing with 75bp read length was carried out using HO flow cell (150 cycle) on the Illumina MiniSeq® instrument.

### NGS Data Analysis

Illumina basespace (https://basespace.illumina.com) bioinformatics pipeline was used for sequencing QC, FASTQ generation, genome assembly, and identification of SARS-CoV-2 variants. Briefly, the Binary Base Call (BCL) raw sequencing files generated by Illumina MiniSeq® sequencing platforms were uploaded to the Illumina basespace online portal and demultiplexed to FASTQ format using the FASTQ Generation (Version: 1.0.0.) application. The raw FASTQ files were trimmed, sorted, and checked for quality (Q>30) using the FASTQ-QC application within the basespace. QC passed FASTQ files were aligned against the SARS-CoV-2 reference genome (NCBI RefSeq NC_045512.2) using Bio-IT Processor (Version: 0×04261818). Then, DRAGEN COVID Lineage (Version: 3.5.4) application in basespace was used for generating a single consensus FASTQ file, for each sample sequenced, on a single flow-cell and variant detection. Finally, single consensus FASTQ was also analyzed for lineage assignment using the web version of Phylogenetic Assignment of Named Global Outbreak Lineages (PANGOLIN) software (https://pangolin.cog-uk.io). Only the consensus variants identified by both applications were used for further analysis.

### Phylogenic Analysis

The FASTQ sequence file was analyzed and visualized for evolutionary relationships through the open-source toolkit Nextstrain (https://clades.nextstrain.org/). GSAID database for global SARS-CoV-2 sequence analysis, available from the Nexstrain server was used to retrieve representative variant sequences.^1^ The NCBI databank was used to retrieve the original Wuhan strain SARS-CoV-2 sequence. All the individual consensus genome sequence files were aligned by using Clustal-W multiple sequence alignment tool.^2^ The phylogenetic analysis was carried out utilizing the Clustal omega server and the phylogenetic tree was constructed using Mega X tool^3^ with default parameters of maximum likelihood method.

The further analysis aimed at investigating the conservation of spike protein in reference sequences vs clinical strains of SARS-CoV-2 from our study using bioinformatics tools. The protein sequences for different ORFs were determined by either annotation by IBM Functional Genomics Platform.^4^ T-COFFEE and PRALINE software^5,6^ were used for the alignment of spike proteins from different isolates and mutation position analysis.

### COVID-19 Lineage assignment using Q-PCR

Commercially available assays from two vendors (GT Molecular [Colorado USA], and Thermo Fisher Scientific [Massachusetts, USA]) were evaluated for detection of known variants, and results were compared to the NGS-based variant detection of the same samples (Table-3). Assays from GT Molecular detected 7 spike protein mutations (N501Y, Del69-70, E484K, K417N, K417T, L452R, T478K).

Assays were provided in two different kits containing the variant-specific reference standard and mutation-specific primer-probe. Amplifications were performed according to the manufacturer’s instructions in four separate master mix preparations as described in Table-3. Briefly, RNA was reverse transcribed for 10 minutes at 53°C followed by enzyme activation 2 minutes at 95°C, and 40 cycles of 15 seconds at 95°C for Denaturation and 60 seconds at 52°C for Annealing/Extension. Reactions were performed by using qScript 1-Step Virus ToughMix (Quantabio Inc, Beverly, MA USA) on LightCycler® 480 System (Roche).

Two TaqMan assays from Thermo Fisher Scientific targeting two spike protein mutations (L452R and P681R) were performed according to the manufacturer’s instructions on LightCycler® 480 System (Roche). The delta variant classifying mutation (L452R) was used for the final classification of the Delta variant across all the Q-PCR-based methods evaluated in this study. Whole genome sequencing followed PANGOLIN classification was used as the gold standard for the final variant classification and validation of the Q-PCR-based variant detection methods.

